# SARS-CoV-2 Spike Affinity and Dynamics Exclude the Strict Requirement of an Intermediate Host

**DOI:** 10.1101/2021.08.11.455960

**Authors:** Matteo Castelli, Luigi Scietti, Nicola Clementi, Mattia Cavallaro, Silvia Faravelli, Alberta Pinnola, Elena Criscuolo, Roberta Antonia Diotti, Massimo Clementi, Federico Forneris, Nicasio Mancini

**Author notes:** These authors contributed equally to this work (Matteo Castelli, Luigi Scietti). These authors jointly supervised this work (Federico Forneris, Nicasio Mancini).

## Abstract

SARS-CoV-2 proximal origin is still unclear, limiting the possibility of foreseeing other spillover events with pandemic potential. Here we propose an evolutionary model based on the thorough dissection of SARS-CoV-2 and RaTG13 – the closest bat relative – spike dynamics, kinetics and binding to ACE2. Our results indicate that both spikes share nearly identical, high affinities for *Rhinolophus affinis* bat and human ACE2, pointing out to negligible species barriers directly related to receptor binding. Also, SARS-CoV-2 spike shows a higher degree of dynamics and kinetics optimization that favors ACE2 engagement. Therefore, we devise an affinity-independent evolutionary process that likely took place in *R. affinis* bats and limits the eventual involvement of other animal species in initiating the pandemic to the role of vector.

## Introduction

The *Coronaviridae* family comprises seven species of human interest; four are endemic and highly adapted to humans (HCoV-229E, HCoV-NL63, HCoV-OC43 and HCoV-HKU1), two epidemic (MERS-CoV and SARS-CoV-1) and one pandemic (SARS-CoV-2). Except for HCoV-OC43 and HCoV-HKU1, their ancestral origin can be traced back to coronaviruses (CoV) infecting bats, the main natural host reservoir of α- and β-coronavirus genera (*1–3*). Epidemic CoVs are characterized by low inter-human transmission and high fatality rate, indicative of a zoonotic infection and a sub-optimal adaptation to humans, and gained the ability to infect humans following adaptation in a putative intermediate host, although with different evolutionary trajectories (*4*). While MERS-CoV ancestors stably adapted to dromedary camels decades ago diverging from bat MERS-related viruses, SARS-CoV-1 direct ancestor seems to have transiently jumped from horseshoe bats (*Rhinolophus* spp.) to palm civets and/or raccoon dogs, accumulating a few mutations that incidentally increased its ability to infect humans.

SARS-CoV-2 was first identified in the city of Wuhan in December 2019 and rapidly spread worldwide due to a high inter-human transmission rate and a relevant percentage of asymptomatic and paucisymptomatic infections (5, 6). Phylogenetic analysis identified SARS-CoV-2 as a member of a novel clade in the *Sarbecovirus* lineage that also comprises viruses identified in Southeast Asian pangolins (*Manis javanica* and *Manis pentadactila*) and horseshoe bats (*7–9*). Among them, RaTG13, collected in 2013 from a *R. affinis* specimen in China’s Yunnan province, shows the highest homology to SARS-CoV-2 both genome-wide and at the spike gene level, thus supporting its bat origin (*5*, *10*). In analogy with SARS-CoV-1 and several bat SARS-related (SARSr) viruses, both RaTG13 and SARS-CoV-2 engage the host angiotensin-converting enzyme 2 (ACE2) to mediate cell entry despite the high sequence divergence at the receptor binding domain (RBD) (*11*, *12*).

In principle, a virus spillover probability is directly proportional to the phylogenetic distance between donor and recipient species, and adaptation in an intermediate host may serve to lower the species barrier. Compared to other viruses, CoVs can jump among host species with relative ease and the major tropism determinant is represented by the spike ability to mediate entry, in turn mainly dependent on the host receptor orthologues conservation. Under this perspective, ACE2 differences between humans and *R. affinis* argue against a direct spillover event. Indeed, albeit still competent, RaTG13 spike binds to human ACE2 (hACE2) and mediates pseudotyped virus entry at a lower extent than SARS-CoV-2 (*12, 13*). While this favors the hypothesis of an intermediate host, no evidence of it have emerged so far, leaving several unanswered questions on the evolutionary path followed by SARS-CoV-2 and posing major concerns on the possible emergence of related viruses with pandemic potential (14).

To trace SARS-CoV-2 evolutionary trajectory, we used a combination of surface plasmon resonance (SPR), X-ray crystallography and molecular dynamics (MD) simulation-based techniques to characterize the functional features of RaTG13 and SARS-CoV-2 spikes. Despite sequence divergence, we found that both RBDs engage hACE2 with nearly identical binding mode and affinity. Furthermore, we measured comparable affinities in the nanomolar range also for *R. affinis* ACE2 (*affiACE2*). At the spike level, SARS-CoV-2 is significantly more optimized to expose the RBD in the conformation competent to ACE2 binding and mutations in all domains contribute to it. Taken together, our results point out to an evolutionary process that regarded exclusively the spike dynamics and kinetics through the fine-tuning of the pre-fusion states metastability. Also, RaTG13 and SARS-CoV-2 RBD promiscuity rules out the requirement of an intermediate host to lower the species barrier.

## Results

### The binding of RaTG13 and SARS-CoV-2 RBD to affiACE2 and hACE2 is nearly identical

Several bat SARSr viruses have the ability to recognize hACE2, surprisingly often at higher affinity than bat ACE2, without prior adaptation (15). The marked sequence differences between RaTG13/SARS-CoV-2 RBDs (89.2% amino acid identity) and human/*R. affinis* ACE2 (80.7% identity), and in particular those found at the RBD-ACE2 interface (Figure 1a), suggest high species barriers to efficient binding and therefore the need of adaptation in an intermediate host. To verify this hypothesis, we first measured the affinity of RaTG13 and SARS-CoV-2 RBDs for hACE2 by SPR. Surprisingly, we found that both RBDs bind to hACE2 with Kd in the nanomolar range (21 and 41 nM, respectively) (Figure 2a). We confirmed SPR measurements through *in silico* free energy calculations (Table S1) and, prompted by these observations, we attempted crystallization of the RaTG13 RBD/hACE2 complex, obtaining crystals suitable for diffraction experiments in two distinct crystal forms (Table S2). After structure determination using molecular replacement, crystal form 1 yielded a 4.5 Å resolution structure of the RaTG13 RBD/hACE2 complex (Figure 2b) showing identical crystal packing assembly to that of the previously determined SARS-CoV-2 RBD/hACE2 complex, whereas crystal form 2 could be solved at 6.5 Å resolution, revealing a different crystal packing assembly (Figure S1) (*12*). Both RaTG13 RBD/hACE2 structures are superimposable to SARS-CoV-2 RBD/hACE2, with minor adjustments associated to the amino acid differences. Therefore, sequence differences between RaTG13 and SARS-CoV-2 RBD do not affect the binding mode nor the affinity for hACE2.

**Figure 1.**
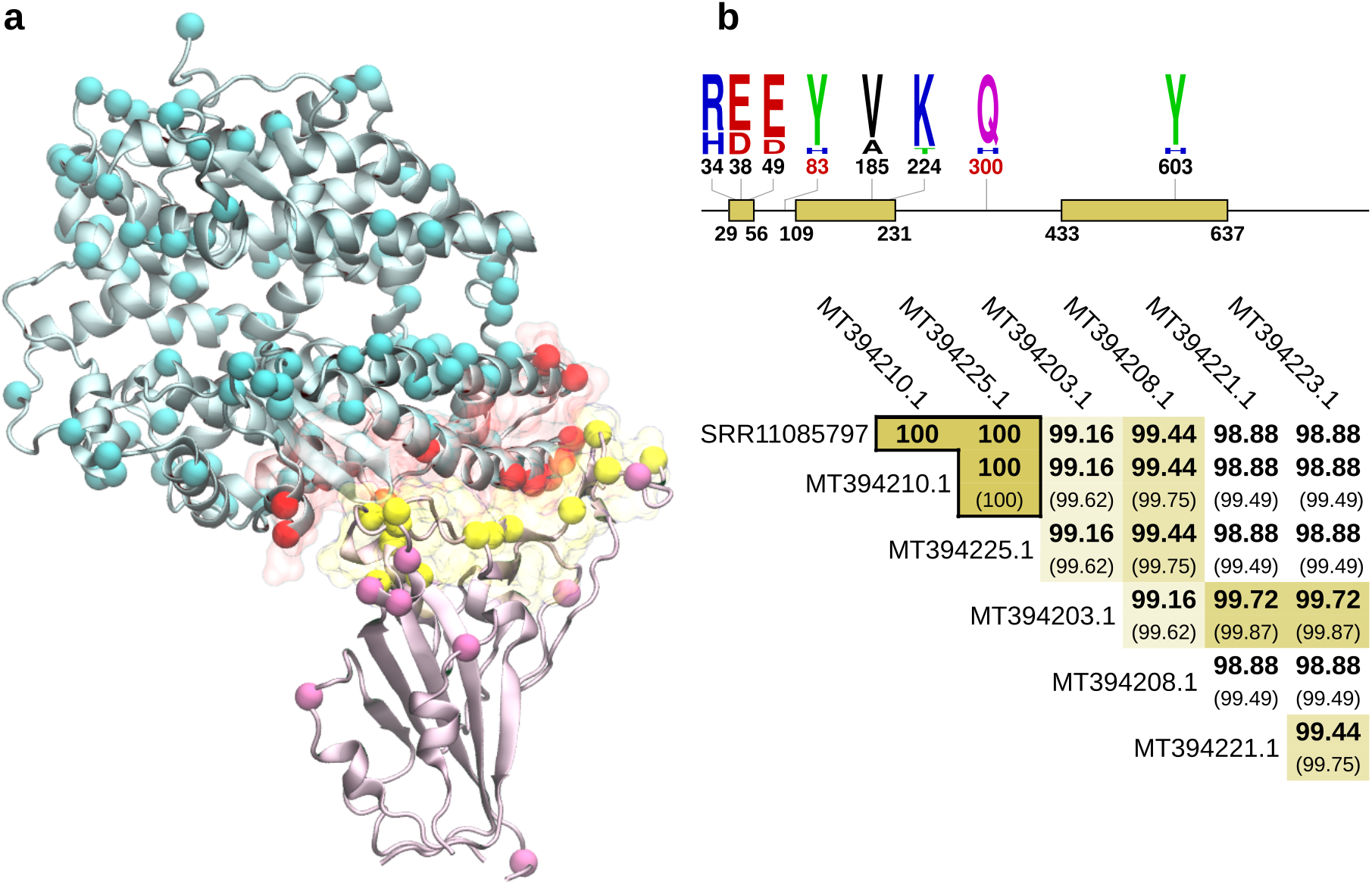
RBD/ACE2 comparison and *affiACE2* allele identification. **(a)** Amino acid differences of RaTG13/SARS-CoV-2 RBD and *affi*ACE2/hACE2. Proteins are depicted in ribbon, with the RBD/ACE2 binding interface at 8 Å in yellow and red transparent surface, respectively. Amino acid differences are depicted as spheres colored according to the distance from the binding interface: RBD and ACE2 mutations below 8 Å are in yellow and red, mutations above 8 Å in purple and cyan, respectively. **(b)** Identification of *affi*ACE2 allele associated to RaTG13. The regions of *affi*ACE2 mRNA covered by SRA reads from dataset SRR11085797 are depicted as yellow bars in the upper panel. Polymorphic sites and relative frequencies were generated with WebLogo (16). Sites not covered by SRA reads are reported in red. Amino acid identity percentage considering covered regions (in bold) or the entire deposited sequences (in brackets) are reported in the lower panel. Identical sequences were collapsed into a single representative. The full comparison of all deposited *affi*ACE2 sequences is reported in Figure S2.

**Figure 2.**
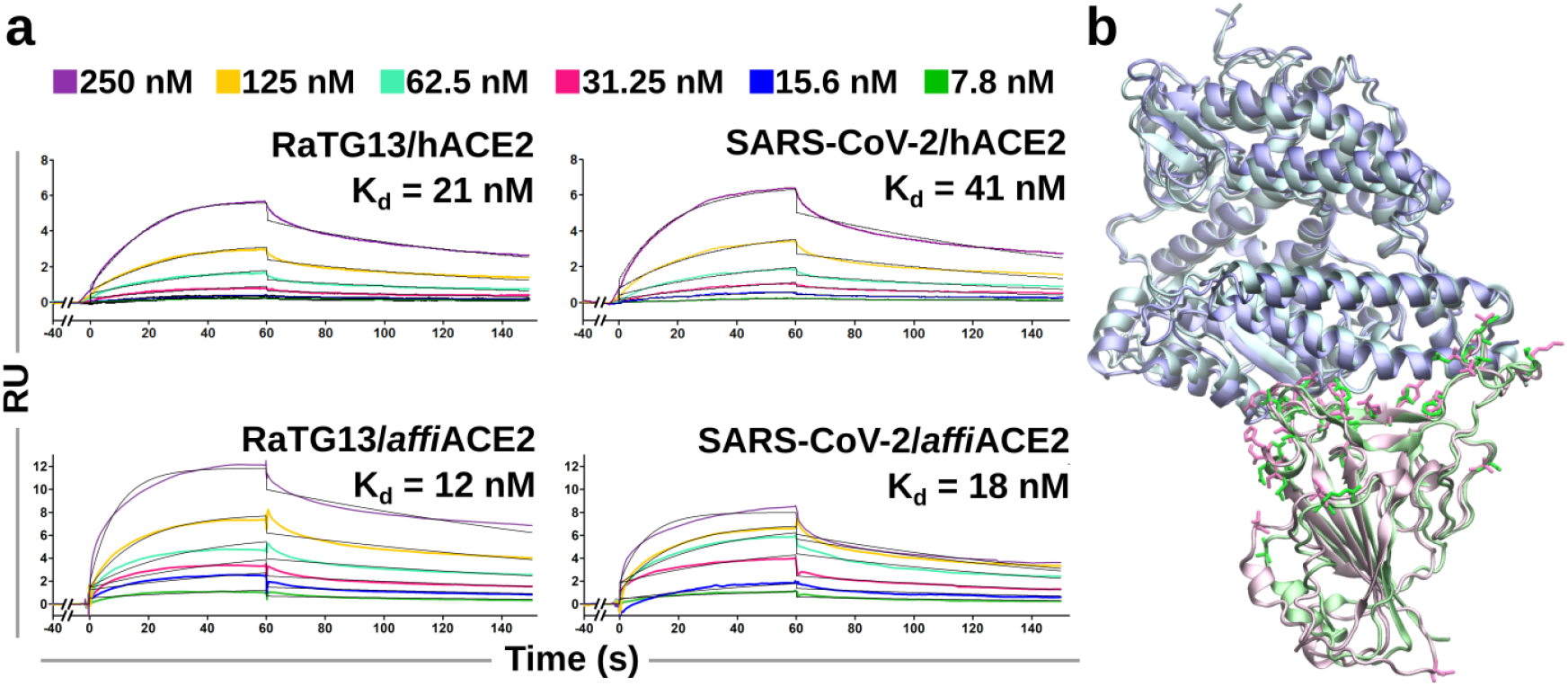
RBD/ACE2 binding affinity and structure. **(a)** Surface plasmon resonance measurements. Blank subtracted sensograms (black curves) of the RaTG13 and SARS-CoV-2 RBDs on immobilized hACE2 and *affi*ACE2. A 1:1 binding model was used for data fitting. Shown data are the mean of four replicates. **(b)** Structure comparison of SARS-CoV-2 (PDB ID: 6M17) and RaTG13 (reported here) RBD/hACE2 complexes. Whole structures are depicted in ribbon, hACE2, RaTG13 RBD and SARS-CoV-2 RBD are colored in shades of blue, pink and green, respectively. The side chain of RaTG13/SARS-CoV-2 mutations are reported in licorice.

To further characterize the evolutionary trajectory of SARS-CoV-2, we next measured RBDs affinity for *affi*ACE2. Several *affi*ACE2 sequences were recently deposited and show moderate variability, with eight polymorphic positions (*15*). We mined the original raw sequencing dataset RaTG13 was identified from and uniquely determined that the specific *R. affinis* specimen carried a minority allele, characterized by the H34, D38 – both lying at the RBD/ACE2 interface – and A185 polymorphisms (Figure 1b). SPR measurements show RaTG13 and SARS-CoV-2 RBD affinities for *affi*ACE2 almost identical to hACE2 (12 and 18 nM, respectively), in agreement with *in silico* calculations (Figure 2A and Table S1). Thus, in terms of RBD affinity, the species barrier between *R. affinis* bats and humans is negligible for both RaTG13 and SARS-CoV-2, strongly supporting the possibility of a direct species jump from bats to humans of SARS-CoV-2 and related viruses. As a consequence, SARS-CoV-2 might have directly evolved in *R. affini*s bats.

### RaTG13 Spike Dynamics is Suboptimal for ACE2 Engagement

Our results indicate that the affinity of RaTG13 and SARS-CoV-2 RBD for hACE2 is equivalent. However, when the entire spike is considered, SARS-CoV-2 is a significantly better hACE2 binder and mediates pseudotyped virus entry more efficiently (12, 13). RaTG13 spike cryoEM structures show exclusively the closed state, while SARS-CoV-2 uncleaved, S0 form is found also in the 1-up state (one RBD exposed and two closed), suggesting that different functional properties might be related to the spike propensity of exposing the RBD for ACE2 engagement. To verify this hypothesis, we performed full-atom, unbiased MD simulations of the entire ectodomains in the closed and 1-up states for both SARS-CoV-2 and RaTG13 and monitored their RBDs geometry – opening (Ψ) and rotation (Φ) angles, as defined in Figure 3a – and interactions. The closed protomers are stable in all simulated systems, but RaTG13 spike acquires a more compact structure, as evidenced by the closed RBDs lower rotation angle and RBD-RBD stronger contacts and correlated movements in both state (Figure 3b-c and S4). These conformational differences are far more marked in the open RBD of the 1-up states. Indeed, while SARS-CoV-2 S0 shows an RBD opening comparable to the starting cryoEM structure, RaTG13 rapidly stabilizes in an intermediate conformation between the open and closed states (Figure 3b and S3), losing most of the interactions and correlated motions between the open and clockwise closed RBDs (Figure 3c and S4). To test whether RaTG13 RBD suboptimal exposure is sufficient to allow the binding to ACE2, we reconstructed the spike/hACE2 complex and found it to be stable over a 200 ns MD simulation, thus confirming RaTG13 competence for receptor engagement in this conformation (Figure S5). Nonetheless, this intermediate aperture provide an explanation of the stark difference in affinity for hACE2 between RaTG13 spike and RBD and its lower functionality compared to SARS-CoV-2.

**Figure 3.**
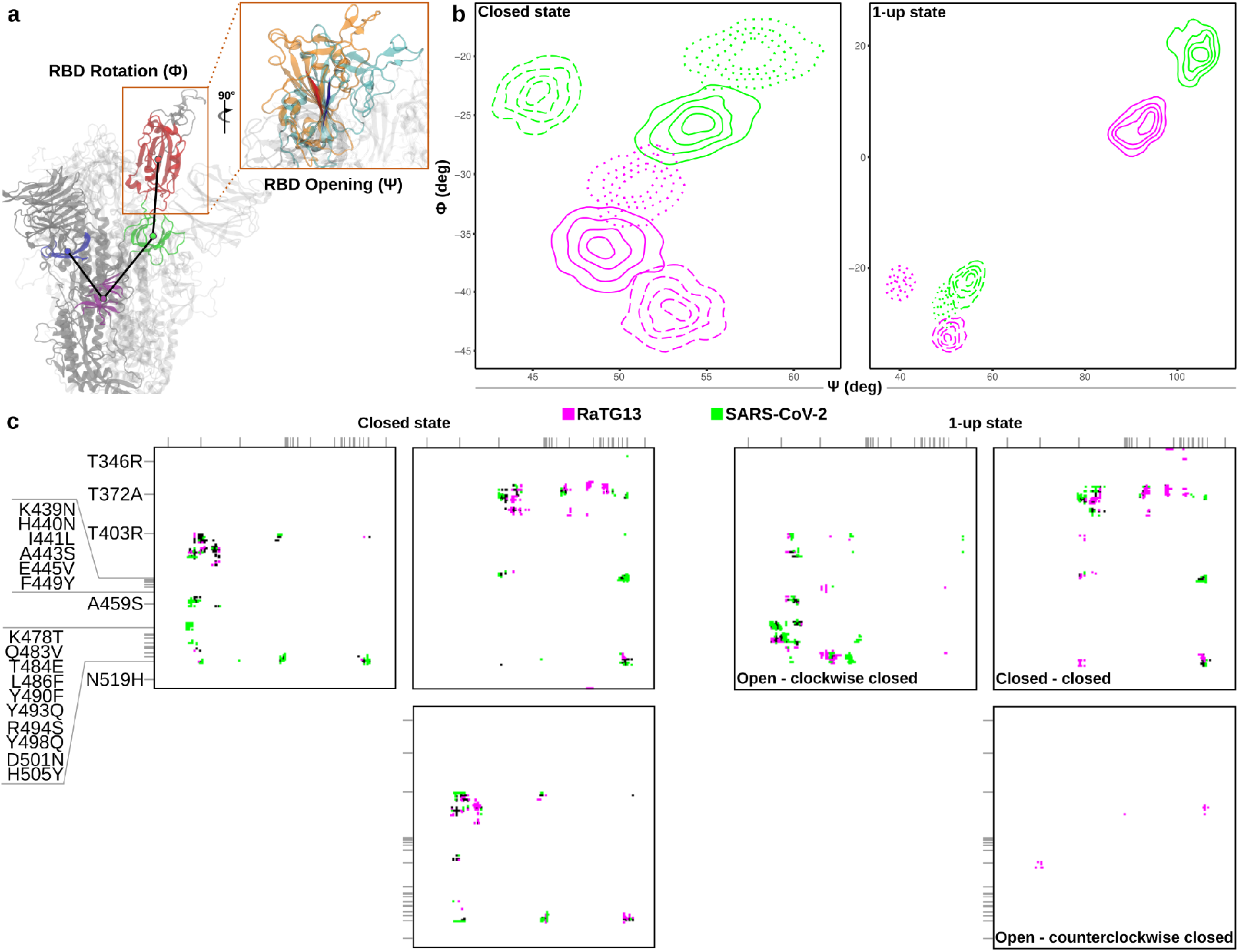
RBD interactions and geometry. **(a)** Definition of the dihedral and angle representative of RBD opening (Ψ) and rotation (Φ), respectively. **(b)** Contour plot of RBDs angles of uncleaved SARS-CoV-2 and RaTG13 spikes in the closed and 1-up states calculated along MD simulations. In the 1-up state graph, solid lines represent the open RBDs, dashed and dotted lines the closed RBDs. **(c)** RBD-RBD avarage contact map calculated along MD simulations. Purple and green black spots indicates contacts specific for RaTG13 or SARS-CoV-2, black spots indicate shared contacts. RaTG13/SARS-CoV-2 mutations are reported on the axes.

To further explore the dynamic properties of RaTG13 and SARS-CoV-2 spike, we next performed targeted MD simulations (TMD) with a linearly increasing bias over 200 ns to drive the closed-to-1-up and 1-up-to-closed transitions using the conformations previously identified in our unbiased MD simulations. The 1-up-to-closed transition is similar between RaTG13 and SARS-CoV-2, as both follow a biphasic kinetics with an initial lag phase – related to the displacement of the counterclockwise protomer N-terminal domain (NTD) and RBD required to accommodate the closure movement – followed by a rapid RBD closure (Figure 4). Conversely, the closed-to-1up transitions follow markedly different paths: as a consequence of the closed RBDs tight interactions, RaTG13 opening is slow and geometrically almost linear, while SARS-CoV-2 S0 is faster and follows a biphasic transition. Its kinetics presents a preparatory initial phase where the RBD motion changes slowly and a subsequent abrupt switch to the open state, indicating that the spike reached a conformation where the RBD is free to move. Altogether, the mutations from RaTG13 to SARS-CoV-2 affect the entire spike pre-fusion states metastability and, by optimizing the RBDs angle of exposure and transitions kinetics, they increase infectivity regardless of the direct affinity for ACE2.

**Figure 4.**
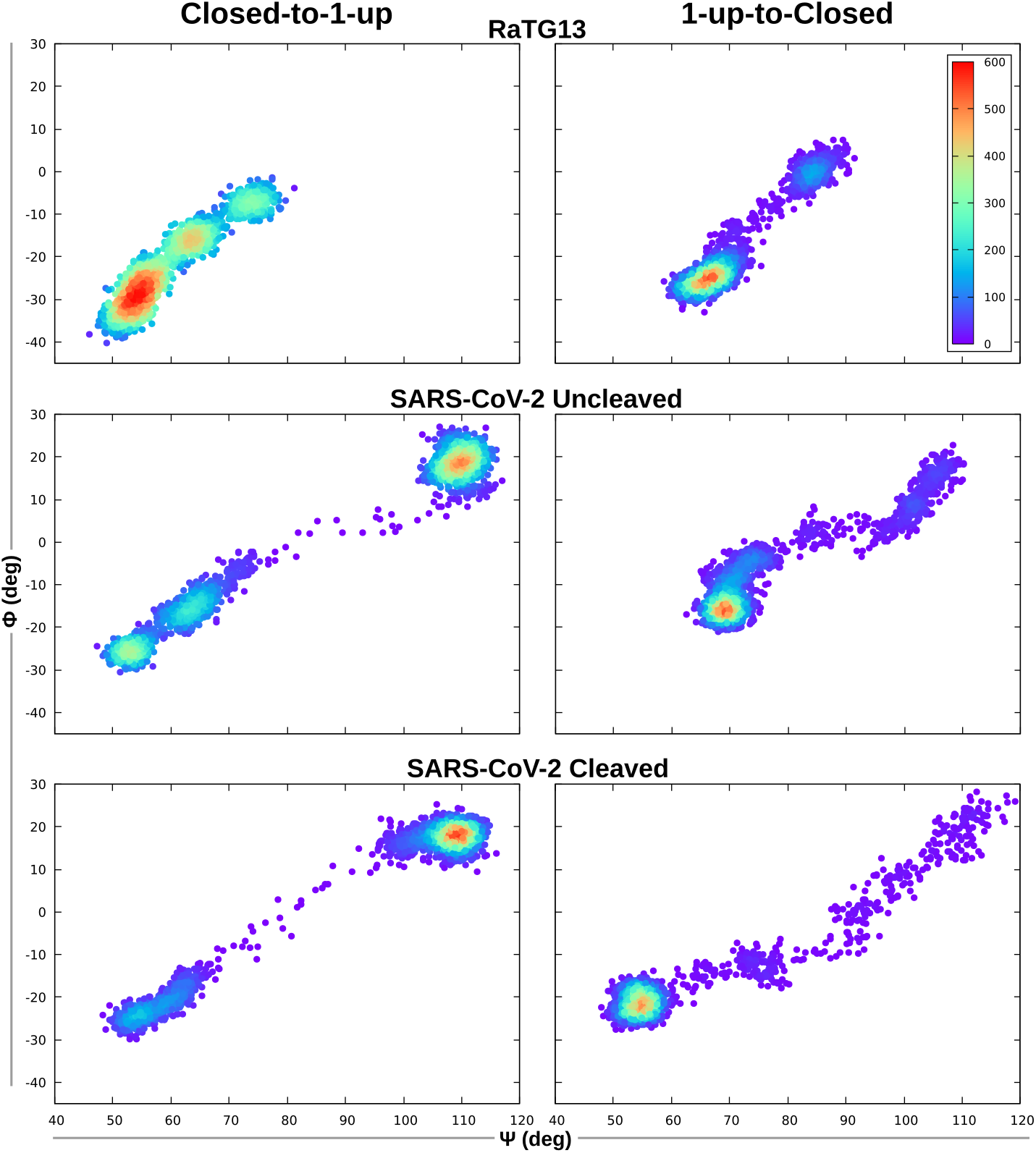
RBD transitions. TMD simulations of RBD opening/closing. The transitions are monitored considering the Ψ and Φ angles of the transitioning RBD and plotted as a function of density.

### RaTG13 Spike Mutations to SARS-CoV-2 Cooperatively Regulate RBD Aperture

Besides the 21 mutations in the RBD, RaTG13 and SARS-CoV-2 spikes differ on four positions in the NTD, one in each subdomain 1 and 2 (SD1 and SD2, respectively), two in S2 (of which only S1125N is comprised in the cryoEM structures and our systems) and the furin cleavage site, a four-residue insertion (^681^PRRA^684^) at the S1/S2 junction (Figure 5a). Hence, the dramatic differences between RaTG13 and SARS-CoV-2 spike dynamics may be due to the local effect of RBD mutations, the allosteric effects of distal insertion/mutations in other domains or the furin cleavage during spike biogenesis. A direct involvement of the RBD is supported by the negligible differences in affinity and by the fact that several mutated residues lie either at the closed RBD-RBD interface (position 372, 403, 439, 440, 498, 501 and 505) that stabilizes RaTG13 or between the open RBD and that in the clockwise closed protomer (position 372, 478 and 486) that stabilizes SARS-CoV-2 open RBD (Figure S6). To test this, we introduced SARS-CoV-2 RBD in the RaTG13 spike backbone alone (RaTG13_RBD_), in combination with the furin cleavage site (RaTG13_RBD+PRRA_) or with the mutations in NTD, SD1, SD2 and S2 (SARS-CoV-2_ΔPRRA_). Worth mentioning, SARS-CoV-2_ΔPRRA_ subpopulations naturally occur at very low frequency *in vitro* and in patients, demonstrating its residual functionality but also a markedly lower fitness (17–19). The 1-up state of RaTG13_RBD_ and SARS-CoV-2_ΔPRRA_ shows large RBD opening (Ψ ~130° and ~120°, respectively) and rotation angles (Φ ~50° and ~60°, respectively) (Figure 5b). Similar Ψ values are found in SARS-CoV-2 cryoEM structures, but Φ values are always below ~35°, suggesting a functional limit to RBD rotation (*20*, *21*). To confirm this hypothesis, we applied a steered molecular dynamics (SMD) protocol to SARS-CoV-2 3-up spike fully engaged with ACE2 to drive all RBDs to Φ = 60°. Reaching the target rotation angle indeed also causes large Ψ increases (up to 165°) and dramatic S1 rearrangements (Figure S7), thus relating SARS-CoV-2_ΔPRRA_ low fitness to prevented furin cleavage and disrupted spike conformation. Conversely, RaTG13_RBD+PRRA_ has an opening angle comparable to SARS-CoV-2 S0 and a remarkably low rotation (Φ ~-10°). Since we recently characterized a SARS-CoV-2 clinical isolate with analogous Φ angle and driving the 3-up state to this Φ value in SMD does not alter the spike global conformation (Figure S7), we can speculate that RaTG13_RBD+PRRA_ geometry is functional (*22*). Therefore, we next analyzed its closed-to-1up transition in TMD and identified a biphasic kinetics similar to SARS-CoV-2 S0, although smoother and slower (Figure S8). The hybrid systems involving SARS-CoV-2 RBD confirm that the sequence of this domain dominates the spike dynamics and kinetics. However, the inclusion of mutations in other domains also highlights relevant allosteric effects. In support to this, structural inspection of RaTG13_RBD_ and SARS-CoV-2_ΔPRRA_ trajectories shows that the RBD large aperture is stabilized by the N519H RBD mutation, as the histidine stably inserts into a hydrophobic pocket created by residues V130, F168, L229, I231 and I233 (Figure S9) in the counterclockwise protomer NTD, that is conserved in all tested systems.

**Figure 5.**
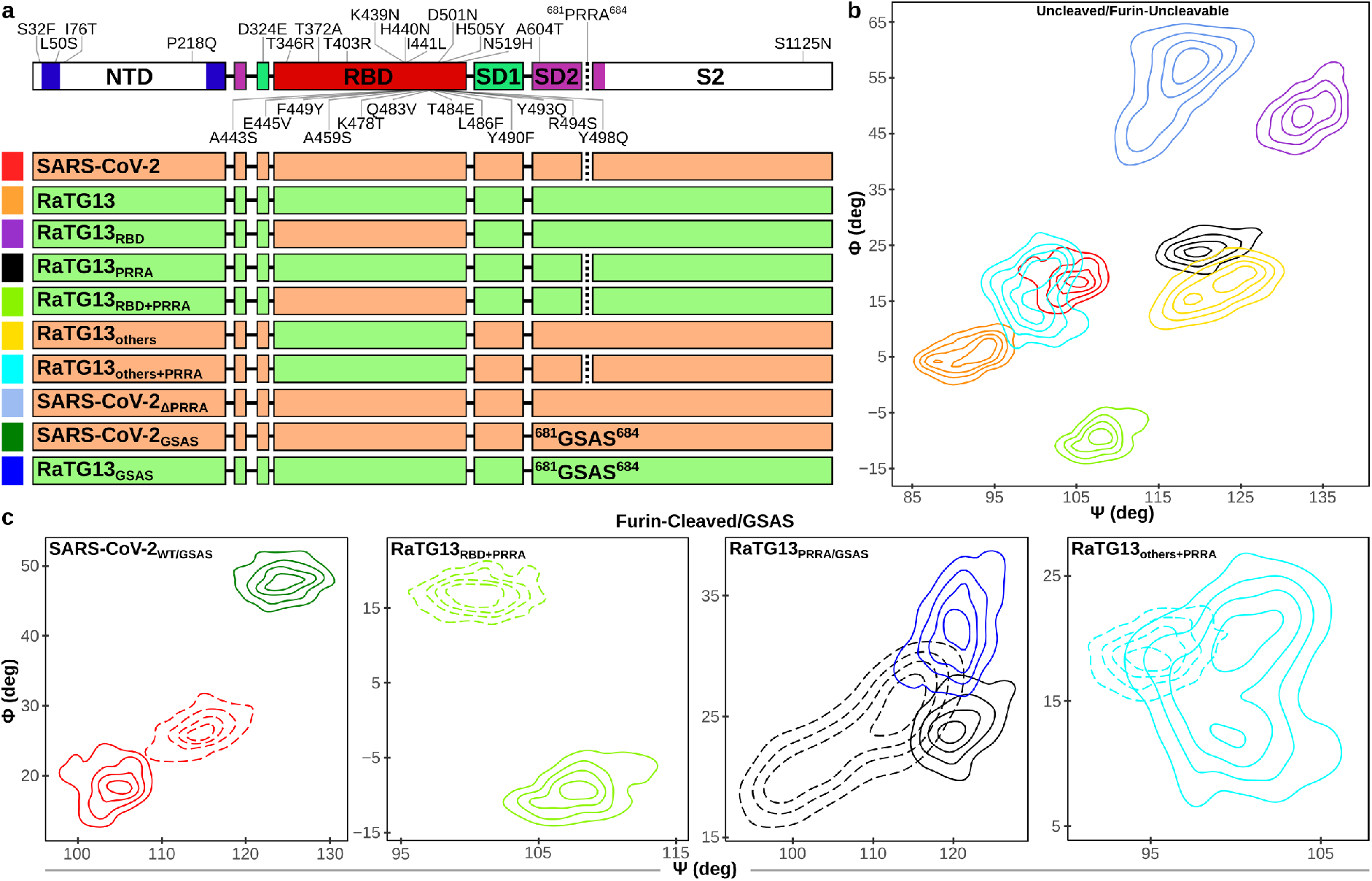
RBD geometry of hybrid spikes. **(a)** Scheme reporting all mutations/insertions from RaTG13 to SARS-CoV-2. The spike is divided into the relevant domains. Color code is the same as in Figure 2a. The green and orange bars below indicate the domains that were swapped among the different hybrid systems, color boxes on the left are the same as in panel B and C. **(b)** Contour plot of the open RBDs Ψ and Φ angles of uncleaved/furin-uncleavable spike hybrid systems. **(c)** Contour plot of the open RBDs Ψ and Φ angles of cleaved and GSAS systems. Solid and dashed lines of the same color represent the uncleaved and cleaved form of the same system, respectively.

To expand each spike domain involvement, we next inserted the cleavage site (RaTG13_PRRA_), mutations in NTD, SD1, SD2 and S2 (RaTG13_others_) or their combination (RaTG13_others+PRRA_) into the RaTG13 spike backbone. RaTG13_PRRA_ and RaTG13_others_ show larger opening angles (Ψ ~120-125°) and a rotation comparable to SARS-CoV-2, while in RaTG13_others+PRRA_ the combination of mutations in all domains but the RBD results in a geometry the closest to the wild-types. Altogether, all domains participate in fine-tuning the spike dynamics through long-range allosteric effects. Our results, together with previous cryoEM structures and functional data, also suggest the existence of functional limits to the RBD geometry. In this scenario, SARS-CoV-2 RBD high propensity to open and rotate may be detrimental in specific spike contexts, outlining a constrained co-evolution of the spike where allosteric effects from the ^681^PRRA^684^ insertion and other domains limit it.

### Furin Cleavage Effects Depend on Residues Located Across the Spike Ectodomain

The furin cleavage site at the S1/S2 junction is found in several unrelated bat β-coronaviruses and a few SARS-CoV-2-related viruses bearing partial furin cleavage site have been recently identified in horseshoe bats, supporting the possibility it arose along the evolution in *R. affinis* (7, 8, 23). While a selective advantage of incorporating it comes from a spike already primed for fusion during biogenesis, cryoEM structures of SARS-CoV-2 S1/S2, cleaved spike demonstrated also a higher propensity to expose the RBDs than the S0 form. To better explore the molecular details associated to furin cleavage and hypothesize when and how the cleavage site was incorporated, we ran unbiased MD simulations of the cleaved, S1/S2 spike form of all wild-type and hybrid systems bearing the ^681^PRRA^684^ sequence (Figure 5).

Compared to the uncleaved form, SARS-CoV-2 S1/S2 1-up state open RBD is in a more upright position achieved through a balanced increase in opening and rotation angles. This conformation is highly similar to that of the ACE2-bound, cleaved spike structure described by Benton *et al*., demonstrating that ACE2 acts by stabilizing rather than inducing it (20). The transitions simulated in TMD show slower 1-up-to-closed and faster closed-to-1up kinetics compared to the uncleaved form. The latter transition is improved both overall and considering exclusively the switch phase, as confirmed by short TMDs performed starting from the end of the lag phase at different fixed biases (Figure 4 and S10). Besides allowing the virus to fuse directly at the plasma membrane in the presence of the TMPRSS2 host protease, this indicates that furin cleavage improves infectivity by increasing RBD aperture and altering the closed/open balance, thus explaining the shift towards the 1-up state evidenced in cryoEM (13).

We next investigated the dynamic properties of furin-cleavable RaTG13/SARS-CoV-2 hybrid spikes to dissect the impact of furin cleavage in association with other mutations.

Cleaved RaTG13_RBD+PRRA_ shows a slight reduction in the opening angle and a marked increase in the rotation compared to the S0 form, thus not reaching SARS-CoV-2 aperture in either forms. Also, cleaved RaTG13_RBD+PRRA_ closed-to-1-up transition follows the same geometric path of the uncleaved form albeit with a faster kinetics (Figure S8), confirming furin cleavage importance in favoring SARS-CoV-2 RBD exposure. Finally, cleaved, 1-up RaTG13_PRRA_ and RaTG13_others+PRRA_ display only minor differences in RBD aperture compared to their respective S0 forms. Altogether, cleaved hybrid systems point out to the role of NTD, SD1, SD2 and S2 mutations in propagating furin cleavage effects to the RBD and suggest that, in the absence of the prone-to-open SARS-CoV-2 RBD, the impact of furin cleavage on the spike dynamics is limited.

Given the furin cleavage site insertion/abrogation relevance in all tested S0 molecular contexts, we finally verified whether its influence is length- or sequence-dependent by mutating ^681^PRRA^684^ into ^681^GSAS^684^ to construct the RaTG13_GSAS_ and SARS-CoV-2_GSAS_ systems. The former 1-up state shows a RBD opening and rotation highly similar to RaTG13_PRRA_, suggesting that the conformational changes may be induced by the increased SD2 length (Figure 5c). However, SARS-CoV-2_GSAS_ 1-up RBD aperture is the closest to SARS-CoV-2RBD, indicating that both SD2 length and a polybasic sequence are needed to compensate for SARS-CoV-2 RBD tendency to open. Altogether, these results strongly support a progressive onset and a continuous co-evolution of the SD2 domain with the RBD, further optimized by adaptation in all spike domains.

## Discussion

Bats species belonging to different families harbor the highest diversity of α- and β-coronaviruses worldwide and have been identified as the ancestral source of five out of seven CoV species of medical interest (1). Also, CoVs diversity proportionally increases with the number of different bat species co-existing in a single habitat and intra-host receptor variability (24). The tight co-evolution of CoVs and bats is particularly evident when SARSr viruses and their long-term natural host – the horseshoe bat *R. sinicus* – are considered. Indeed, high ACE2 diversity is found exclusively at the RBD-binding interface and different ACE2 alleles sustain to a variable extent bat SARSr RBD binding and entry in a virus species-specific manner, suggesting an intra-host spike evolutionary process driven by the transfer between host subpopulations and subsequent adaptation (15). The same study also reported that the RBD of two of the closest SARS-CoV-1 relatives, RsWIV1 and RsSHC014, bound with a 10-fold higher affinity human than *R. sinicus* ACE2 (10^-7^ and 10^-6^ M, respectively), while SARS-CoV-1 RBD displayed the same trend but with a 1000-fold difference in favor of hACE2 (10^-8^ and 10^-5^ M). This demonstrates that, considering exclusively RBD affinity for the receptor, several bat SARSr viruses could in principle infect humans without prior adaptation, but sustained human-to-human transmission requires a tighter binding to hACE2. In addition, RBD mutations accumulated along the evolution in palm civets and humans have increased the affinity for their ACE2 at the expense of the efficient binding to that of the reservoir species. Worth mentioning, a similar process has been postulated for MERS-CoV as well, despite different receptor usage, host species tropism and permanence in intermediate hosts (25). Altogether, RBD/ACE2 affinity of human and closely related, animal viruses for their respective host receptor efficiently recapitulate their evolutionary trajectory, suggesting this affinity-based strategy can be used to characterize other spillover events and trace back SARS-CoV-2 origin.

Our SPR measurements and *in silico* calculations of the RBD/ACE2 complexes point out to unique binding features representative of an evolutionary trajectory radically different from that of SARS-CoV-1. We note that RaTG13 RBD/hACE2 affinity and structure were recently determined and diverge from our results (49). However, some of the proteins used in that study were produced in insect cells, leading to a different hACE2 glycosylation pattern (N90 and N432 glycosylations are uniquely present in our structure, while N322 is absent) that may be at the basis of the discrepancies between SPR measurements. Our results show that SARS-CoV-2 and RaTG13 RBDs both have affinities for *affi*ACE2 and hACE2 in the low nanomolar range (10^-8^ M) despite sequence divergence. Compared to SARS-CoV-1, SARS-CoV-2 displays similar binding to hACE2 but a 1000-fold higher affinity to its closest relative natural host receptor, while RaTG13, compared to SARSr viruses, has a 100- and 10-fold higher affinity for its natural host receptor and hACE2, respectively. Therefore, considering its RBD affinity for hACE2, RaTG13 has a higher potential of directly crossing the species barrier to humans than previously determined for bat SARSr viruses (*26*). Also, the fact that SARS-CoV-2 RBD is endowed with identical binding features and poorly binds to other *Rhinolophus* spp. ACE2 as recently determined strongly suggests the virus directly evolved in *R. affinis* (27). Altogether, the affinity pattern we identified does not fit into an evolutionary model implying an intermediate host and supports SARS-CoV-2 direct spillover from *R. affinis* bats allowed by its unprecedented affinity for the natural host receptor and the promiscuous binding to hACE2 – and several other orthologues – without adaptation (28–30).

The accumulation of a large number of mutations at the RBD that does not result in affinity changes during permanence in the same host is counter-intuitive from an evolutionary perspective, as does not confer any obvious selective advantage. Nonetheless, SARSr RsSHC014 and RsWIV1 show very similar traits: they are highly homologous genome- and spike-wide but 35 mutations are found at the RBD, they have a stable and negligible RBD net charge despite the selection of many charge-changing mutations and they have the same affinity profile for hACE2 and *R. sinicus* major alleles (15). However, their evolution was prompted by *R. sinicus* ACE2 high variability, while *affi*ACE2 is moderately variable, with only two polymorphic positions at the RBD/ACE2 interface. Thus, whether SARS-CoV-2 ancestors circulated in a mixed *R. affinis* population or stably transferred from one subpopulation to the other, RBD direct affinity improvement is not the evolutionary driving force of SARS-CoV-2 spike. Therefore, we extended our analysis at the whole spike level, performing unbiased and enhanced sampling MD simulations of the spike ectodomains to evaluate their dynamics and kinetics. Our results expand the previous knowledge derived from cryoEM structures of SARS-CoV-2 spike S0 and furin-cleaved forms being increasingly more prone to adopt the 1-up state than RaTG13 (13). We demonstrated that RaTG13 1-up state engages the host receptor with a sub-optimal RBD angle that has direct effects on spike affinity and indirect on avidity through the curtailed transition to the 2-up and 3-up states, thus relating the marked discrepancies between RaTG13 and SARS-CoV-2 spike functionality to global structural features. We found that optimal SARS-CoV-2 RBD aperture in the 1-up state and transition kinetics are both determined by RBD-RBD interprotomer interactions and allosteric effects from distal domains. Indeed, SARS-CoV-2 RBD has a higher propensity to open *per se* through looser RBD-RBD interactions that involve protomers in the closed state and tighter interactions between the open and the closed, clockwise RBDs in the 1-up state. RaTG13 mutation to SARS-CoV-2 of each domain in single always leads to a 1-up RBD larger opening angle, and a fundamental regulatory role is played by the furin cleavage site insertion, that always limits RBD opening and rotation when combined to mutations in other domains. Also, the ^681^PRRA^684^ insertion effects on the spike global dynamics are both length- and sequence-dependent and rely on the correct signal propagation to the RBD that is determined by the spike background, as demonstrated by the different conformational rearrangements induced in cleaved hybrid systems compared to SARS-CoV-2. Altogether, our results point out to a progressive co-evolution of SARS-CoV-2 spike that improved receptor engagement exclusively in a RBD direct affinity-independent fashion through the accumulation of single-point mutations. Given the contribution of each domain to the spike dynamics, the strongest selective pressures were on the RBD: its evolution was driven by conformational optimization and constrained by maintenance of direct RBD/ACE2 affinity and proper configuration of titratable amino acids – a trait common to other SARSr viruses likely linked to the spike exposure to mildly acidic pH during cell egress and stomach transit to cause enteric infection in bats (*31*). As such, a discrete recombination event, previously hypothesized to be at the basis of SARS-CoV-2 RBD acquisition in a RaTG13-like spike molecular context, seems unlikely, as it should have taken place after ^681^PRRA^684^ insertion to counteract SARS-CoV-2 RBD intrinsic tendency to open and rotate (*32*, *33*).

In conclusion, we clearly outlined a unique scenario where a human coronavirus and its closest animal relative share the same high-affinity pattern for both the donor and acceptor species, indicating the absence of species barriers related to receptor direct binding. At the spike level, sequence differences between the two viruses account exclusively for global dynamics and conformational optimization that improved infectivity regardless the host species. These features rule out the requirement of SARS-CoV-2 adaptation in an intermediate host; other animal species, if any, have acted as mere vectors in SARS-CoV-2 transfer from bats to humans due to its broad host tropism. Also, they posit the possibility of other spillover events of SARS-CoV-2-related bat viruses with analogous spike features.

## Materials and methods

### RaTG13-associated affiACE2 allele identification

*R. affinis* ACE2 known coding sequences (GenBank database accession IDs MT394203 to MT394225) were fed to Sequence Read Archive (SRA) Nucleotide BLAST to retrieve ACE2 Illumina reads from the SRA dataset with accession ID SRX7724752. Reads were assembled in contigs with Spades (34).

### Generation of plasmid vectors for recombinant protein production

The pCAGGS plasmids for production of the C-terminal His-tagged RBD (#NR_52310), were obtained from BEIRESOURCES (NY, USA). The designed codon-optimized sequence encoding for RaTG13 RBD and the cDNA for *affiACE2* ectodomain (residue 19-615, based on the sequence deposited on Genbank under the ID MT394225.1) were synthesized by Genewiz with flanking 5’-BamHI and 3’-NotI restriction sites and sub-cloned into pUPE.107.03 plasmid vectors (U-Protein Express B.V., The Netherlands) providing the human cystatin protein signal peptide and C-terminal 6xHis-tag for purification. The cDNA for hACE2 ectodomain was obtained from AddGene (#141185). The sequence was amplified using PCR with oligonucleotides hACE2ecto-Fw (5’-aaaatgatcaTCCACCATTGAGGAACAGGCC-3’) and hACE2ecto_Rv (5’-aaaagcggccgcGTCTGCATATGGACTCCAGTC-3’) and sub-cloned into a pUPE.06.45 expression vector (U-Protein Express B.V., The Netherlands) providing the signal sequence from cystatin followed by a removable (TEV-cleavable) N-terminal 6xHis-Strep-tag.

### Recombinant Protein Expression and Purification

Recombinant hACE2 ectodomain, *affi*ACE2 ectodomain, SARS-CoV-2 RBD, and RaTG13 RBD were produced using HEK293-F cells (Invitrogen) cultivated in suspension using Freestyle medium (Invitrogen) as described in (*35*). Briefly, cells were transfected at a cell density of 1 million mL^-1^ using a mixture of 1 μg of recombinant expression plasmid and 3 μg of polyethyleneimine (PEI; Polysciences, Germany). Cultures were supplemented with 0.6% Primatone RL (Merck) 4 h after transfection. The cell media containing secreted proteins were collected 6 days after transfection by centrifugation at 1000 × g for 15 min. The pH and ionic strength of the filtrated medium were adjusted using concentrated phosphate buffer saline (PBS). Samples were loaded onto a 5 mL His-Trap excel column (Cytiva biosciences) using a peristaltic pump and then eluted with a 0-250mM imidazole gradient using a NGC fast protein liquid chromatography (FPLC) system (Bio-Rad). The eluted samples were subject to immediate concentration with concomitant buffer exchange with fresh PBS to remove imidazole using Amicon centrifugal filters (Merck). For hACE2, the N-terminal His-Strep tag was cleaved by incubating the sample with His-tagged TEV protease for 2 hours at RT, followed by affinity-based sample cleanup using a 5 mL His-Trap excel column (Cytiva biosciences). Quality control during protein purification was carried out using reducing and non-reducing SDS-PAGE analysis and differential scanning fluorimetry (DSF) with a Tycho NT.6 instrument (Nanotemper). All samples were concentrated to 1 mg mL^-1^, flash-frozen in liquid nitrogen and kept at −80 °C until usage. Prior to analysis, the protein samples were thawed and subject to gel filtration using a Superdex 200 10/300 increase column (Cytiva lifesciences) equilibrated with 25 mM HEPES/NaOH, 150 mM NaCl, pH 7.2.

### SPR Binding Analysis

hACE2 or *affi*ACE2 were immobilized on a CMD200M SPR chip (XanTec bioanalytics GmbH) using a mixed solution of 200 mM 1-ethyl-3-(3-dimethylaminopropyl) carbodiimide hydrochloride (EDC) and 50 mM N-hydroxysuccinimide (NHS) in a Biacore T-200 SPR instrument (GE Healthcare), reaching 1000 response units (RU). Reactive groups in excess were blocked with 1 M ethanolamine. The system flow cell 1 was pre-activated and blocked using the same procedure and used as reference. Twofold dilutions of SARS-CoV-2 RBD or RaTG13 RBD with concentration ranging from 250 to 7.8 nM were prepared in the running buffer (25 mM HEPES/NaOH, 150 mM NaCl and 0.05% Tween-20, pH 7.2) and injected using a flow of 50 μl min ^−1^. Analysis was performed using the Biacore evaluation software (GE Healthcare) using a 1:1 affinity model.

### Crystallization of RaTG13 RBD/hACE2

RaTG13 RBD and hACE2 were mixed in a molar ratio of 1:1.3 and further subject to gel filtration using a Superdex 200 10/300 increase column (Cytiva lifesciences) equilibrated with 25 mM HEPES/NaOH, 150 mM NaCl, pH 7.5. hACE2-RaTG13 RBD crystals of two different forms were obtained in sitting drop at 25 °C by mixing 0.5 μL of protein complex concentrated at ~10 mg mL^-1^ with 0.5 μL of reservoir solution composed either of 100 mM Tris/HCl pH 8.5, 20-25% PEG 6000, 100 mM NaCl (crystal form #1) or 0.2-0.25 M sodium thiocyanate, 18-23% PEG 3350, pH 6,9 (crystal form #2). Crystals were harvested using MicroMounts Loops (Mitegen), cryo-protected with the mother liquor supplemented with 20% glycerol, flash-cooled and stored in liquid nitrogen prior to data acquisition.

### Diffraction data collection, structure solution and refinement

Diffraction data from hACE2-RaTG13 RBD crystals were collected at the ID23-1 beamline of the European Synchrotron Radiation Facility (ESRF, Grenoble, FR). Data were integrated using the automatic *XIA2-DIALS* pipeline available at the beamline outstation and scaled with AIMLESS (*36*, *37*). Data collection statistics are summarized in Table S1. Individual search models for hACE2 and SARS-CoV-2 RBD molecules were extracted from PDB 6VW1 using *COOT* and used for molecular replament with PHASER, yielding complete solutions for both crystal forms comprising two hACE2 and two RBD molecules in asymmetric unit (*38*, *39*). Final 3D models were generated using iterations of automatic refinement using low resolution protocols in *phenix.refine* using non crystallographic symmetry restraints as well as external restraints based on individual chains from PDB ID 6VW1, alternated with manual adjustments using *COOT* (*40*). Assessment of final structure quality was carried out with the *Molprobity* server and with the RCSB PDB Validation Server (39, 41). Final refinement statistics are listed in Table S1.

### Unbiased Spike Simulations

Deglycosylated spike ectodomains (residues 14-1146) were completed, reverted to the wildtype when needed or mutated with Modeller using the following templates: PDB ID 6ZGF for RaTG13 closed, PDB ID 6ZGG for all cleaved, 1-up systems, PDB ID 6ZGI for all cleaved, closed systems, PDB ID 6VYB for all uncleaved, 1-up system and PDB ID 6ZGE for all uncleaved, closed systems (42). To reconstruct the structure of RaTG13 spike complexed with hACE2, the final frames of the RaTG13 1-up state and RaTG13 RBD/hACE2 simulations were used as templates in Modeller. All structures were simulated in an orthorhombic TIP3P water box, neutralized with the proper counterions, and parametrized using the all-atom AMBER/parm12SB force field (43). All simulations were performed using the GROMACS 5.1.4 code (44). Periodic boundary conditions in the three axes were applied. Covalent bond length, including hydrogen bonds, was set using the LINCS algorithm, allowing a time-integration step of 2 fs. Constant pressure was imposed using the Parrinello-Rahman barostat with a time constant of 2 ps and a reference pressure of 1 bar, while the constant temperature was maintained using the modified Berendsen thermostat with a time constant of 0.1 ps. Long-range electrostatic interactions were calculated with the particle mesh Ewald method with a real-space cutoff of 12 Å. Each system was minimized with the steepest descent algorithm, equilibrated for 100 ps in an NVT ensemble followed by 100 ps in an NPT ensemble, and then subject to a 200ns simulation at constant temperature (300 K).

RBDs contacts were characterized by calculating inter-protomer dynamic cross-correlation and contact map (*45*). RBDs geometry along the trajectories was measured as RBD rotation (Φ) – calculated as the pseudo-dihedral angle generated by virtually connecting the centers of mass of RBD, SD1, SD2 and NTD domains as already reported by Henderson *et al.* – and RBD opening (Ψ) – calculated as the angle subtended by two planes, one perpendicular to S2 central α-helices and passing through residues 1051 Cα of the three protomers and the other through the Cα of RBD residues 354, 395 and 401 (*46*). Residues were chosen for their low RMSF (root mean square fluctuation) values indicating marked intra-domain stability.

### RBD/ACE2 free energy calculations

We generated by homology modeling with Modeller the 3D atomic structures of the RBD/ACE2 (boundaries at residues 334-527 and 19-615, respectively) complexes as follows (42). The atomic structure with PDB ID: 6M17 was used to extract the SARS-CoV-2 RBD/hACE2 complex and as template of SARS-CoV-2 RBD/affiACE2. The crystal structure of the RaTG13 RBD/hACE2 complex obtained here was used *per se* and as template for RaTG13 RBD/affiACE2. MD simulation parameters were the same as for the spike systems and, at the end of the 200 ns run, each system was restarted to produce 10 independent runs of 10 ns each. Binding free energy was calculated as an average of the 10 independent runs using the MM/GBSA method corrected as in (50).

### Enhanced sampling MD Spike Simulations

Spike closed-to-1-up and 1-up-to-closed transitions were simulated using the TMD protocol implemented in Plumed (*47*). The final frame of each state from the unbiased MD 200ns trajectory was used as the starting and final points for the corresponding transition simulated at 300 K. The transitions were driven by applying a linearly increasing spring constant with κ from 0 to 10000 kJoule/mol/nm over 100000000 steps (200ns) exclusively to the Cα of the transitioning protomer structured domains. RBD rotation was simulated using the SMD protocol implemented in Plumed. SARS-CoV-2 spike structure in the 3-up state fully engaged by hACE2 monomers (PDB ID: 7A98) was minimized and equilibrated as previously described and then the three RBDs were driven to the target Φ angle by applying a constant spring with κ = 10000 kJoule/mol/nm over 100000000 steps (200 ns).

## Acknowledgements

We thank Mr. Matteo De Marco for their assistance in molecular cloning and biophysical analyses, and scientists from ARDIS SRL for useful discussions on SPR data processing. We thank Dr. Stefano Iula for useful discussion on geometric measurements. We thank FSTechnology SpA for providing part of the computational resources. We thank the European Synchrotron Radiation Facility (ESRF) for the provision of synchrotron radiation facilities and for the excellent support provided by the ESRF beamline scientists during remote data collection sessions. The following reagents were produced under HHSN272201400008C and obtained through BEI Resources, NIAID, NIH: Vector pCAGGS Containing the SARS-Related Coronavirus 2, Wuhan-Hu-1 Spike Glycoprotein Gene, NR-52310; Vector pCAGGS Containing the SARS-Related Coronavirus 2, Wuhan-Hu-1 Spike Glycoprotein Receptor Binding Domain (RBD), NR-52309.

## Author contributions

Conceptualization: MCas, LS, NC, FF, NM

Methodology: MCas, LS, MCav, NC, FF, NM

Investigation: MCas, LS, MCav, SF, AP, EC, RAD

Visualization: MCas, LS, MCav, FF

Funding acquisition: FF

Project administration: MCas, LS, FF, NM

Supervision: MCl, FF, NM

Writing – original draft: MCas, LS

Writing – review & editing: MCas, LS, NC, MCl, FF, NM

## Data and materials availability

Coordinates and structure factors for the RaTG13 RBD/hACE2 complex have been deposited in the Protein Data Bank under accession codes 7P8I (crystal form I) and 7P8J (crystal form II).

---

**Figure S1.**
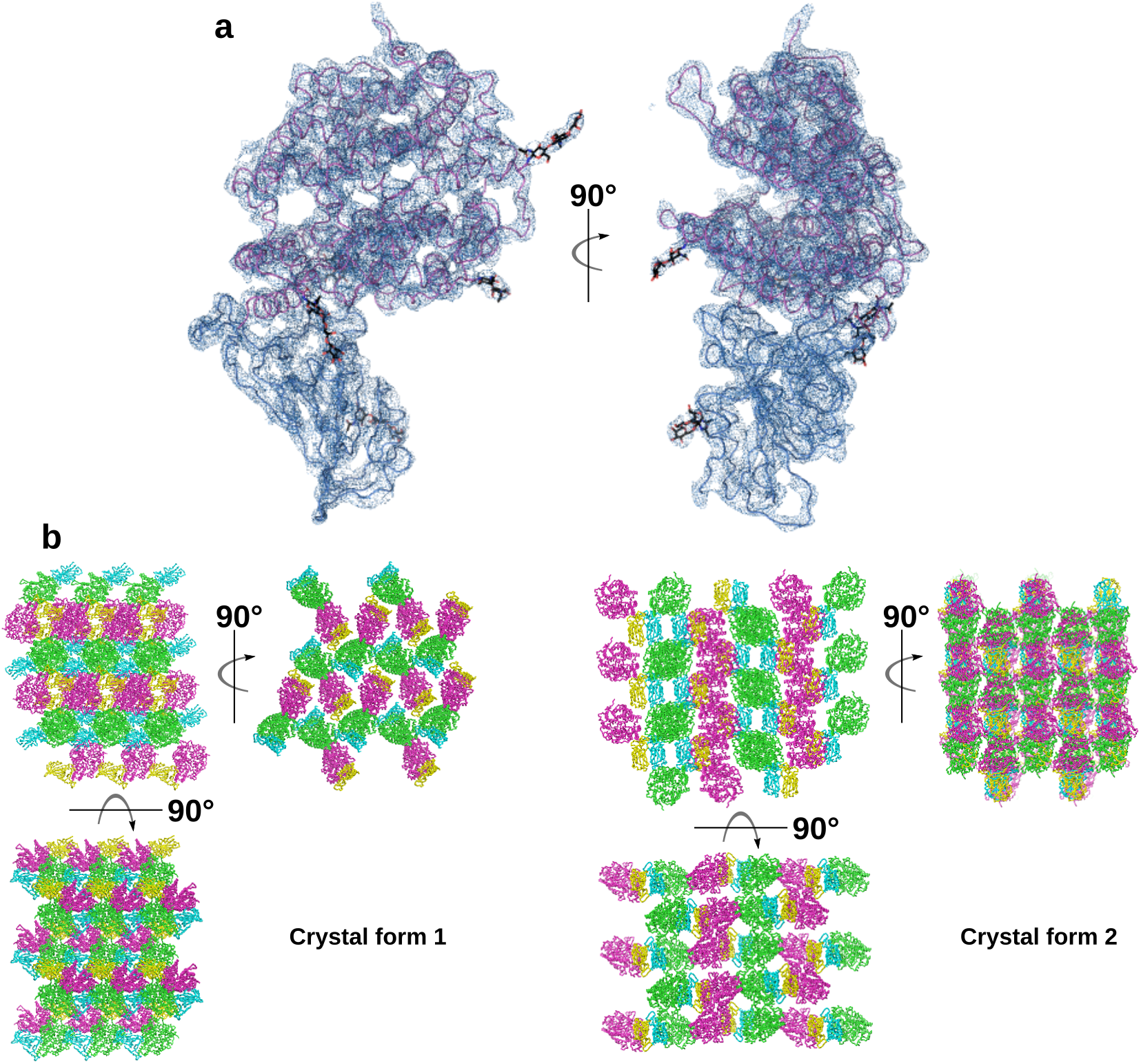
X-ray crystallography of RaTG13 RBD/hACE2 complex. (**a**) Refined 2Fo-Fc electron density maps (contour level 1.1) covering RaTG13 RBD/hACE2 complex (in blue and pink, respectively). (**b**) Crystal packing.

**Figure S2.**
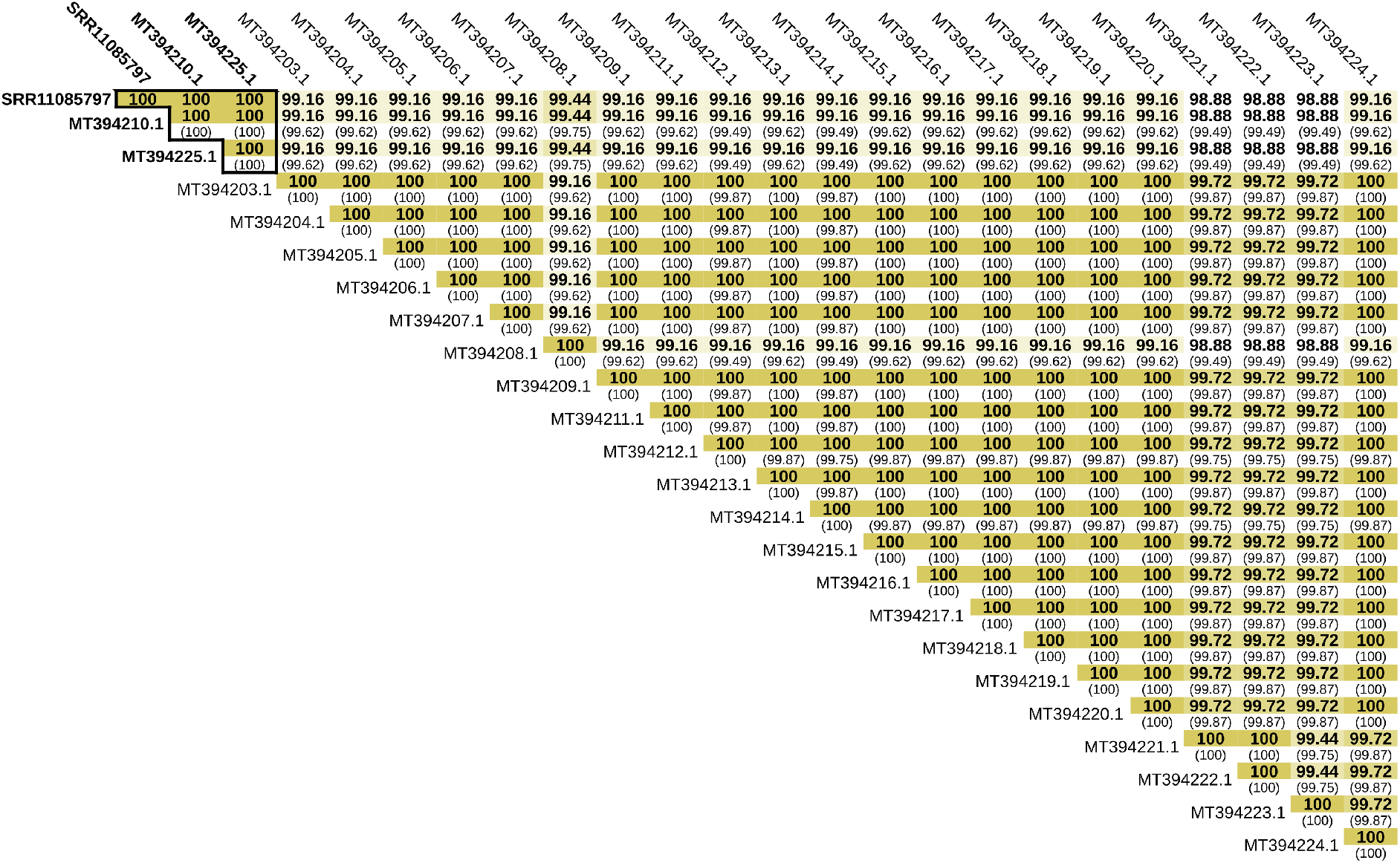
Full *affi*ACE2 alleles comparison. Amino acid identity percentage considering SRA-covered regions (in bold) or the entire deposited sequences (in brackets).

**Figure S3.**
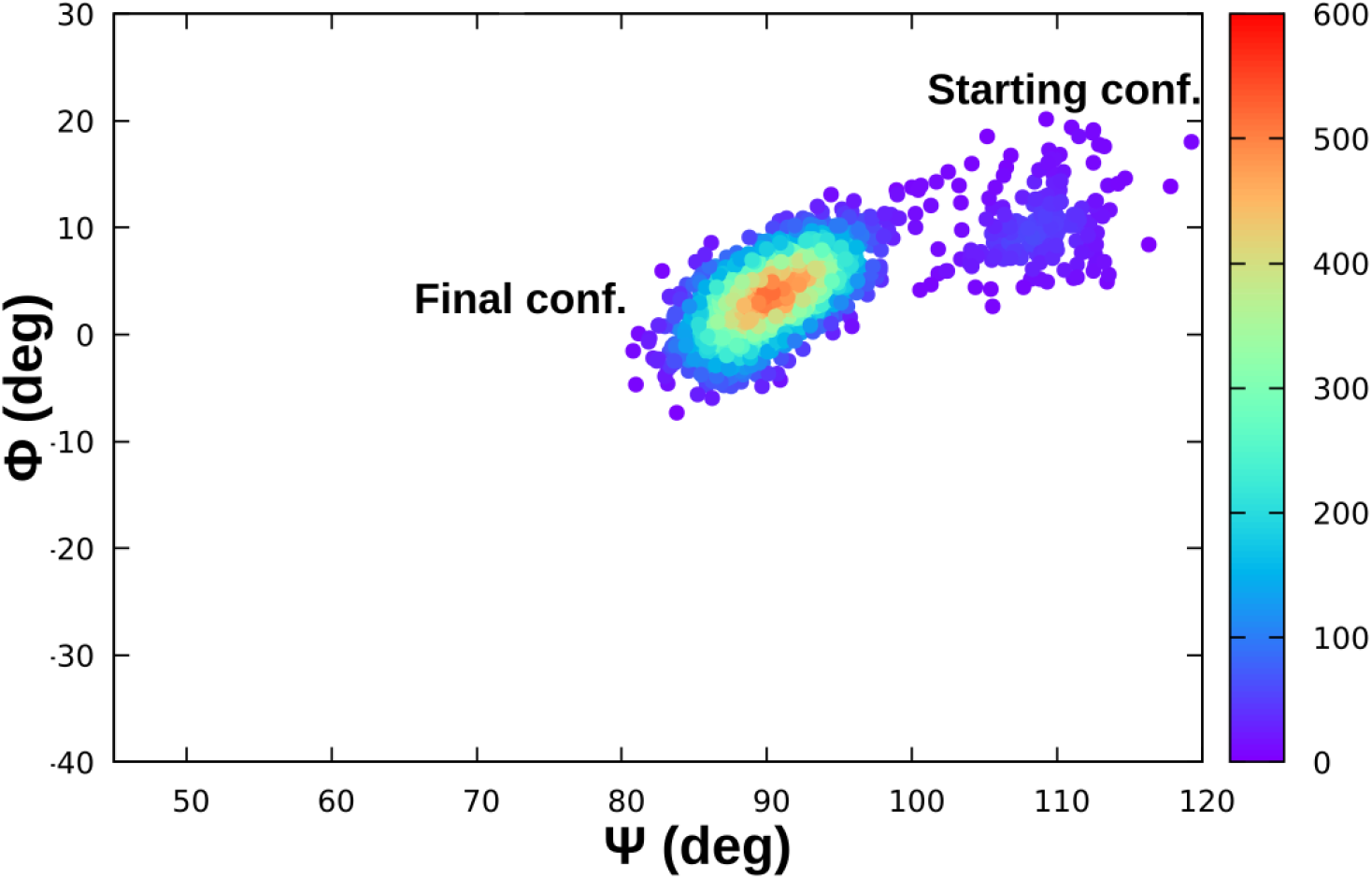
RaTG13 RBD aperture in unbiased MD. Density plot depicting the progressive closure of RaTG13 RBD over 200 ns.

**Figure S4.**
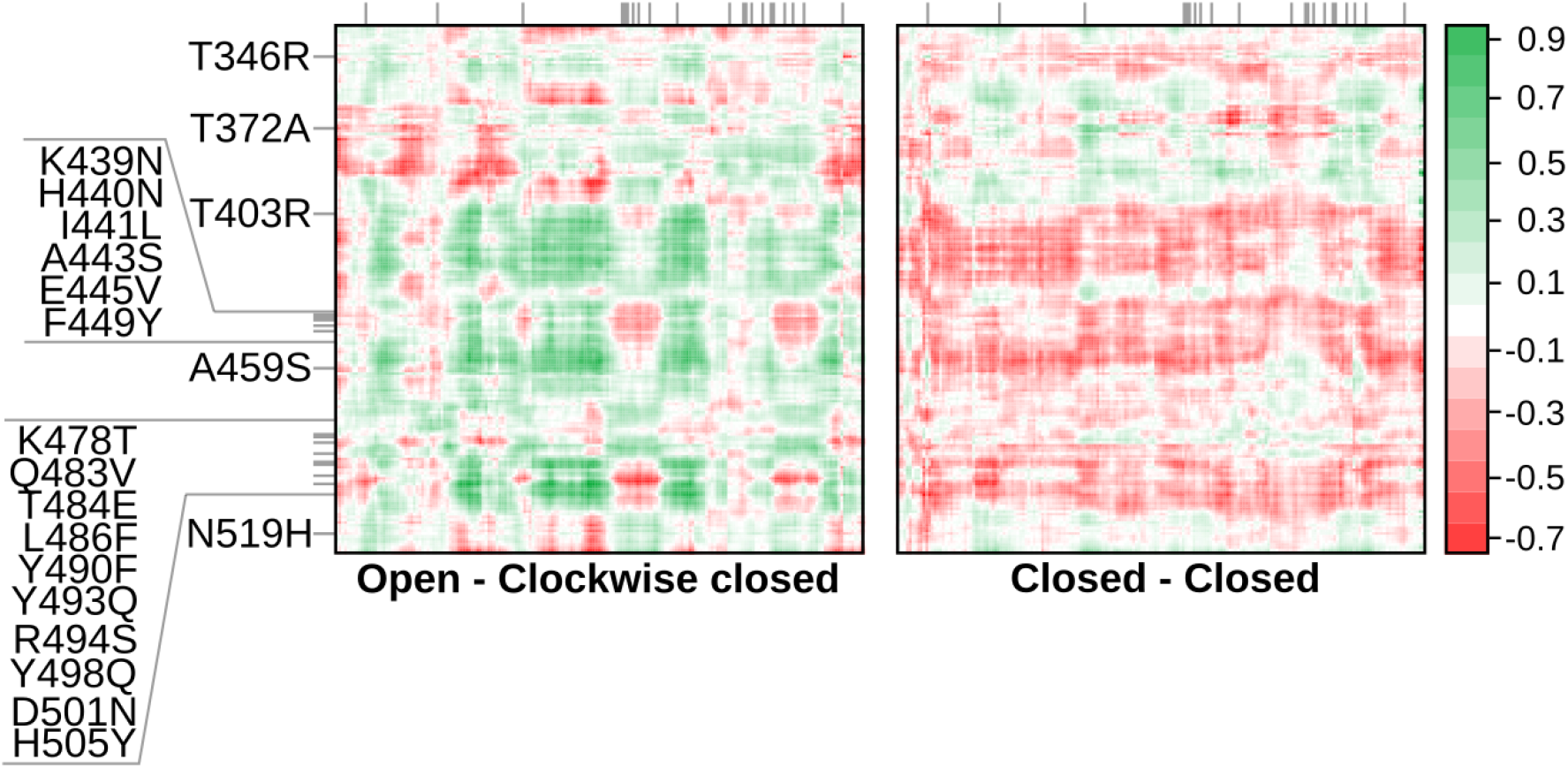
RBD-RBD dynamic cross-correlation. RBD-RBD differential cross-correlation of RaTG13 and S0 SARS-CoV-2 in the 1-up state. Positive values indicate increased cross-correlations in SARS-CoV-2 compared to RaTG13.

**Figure S5.**
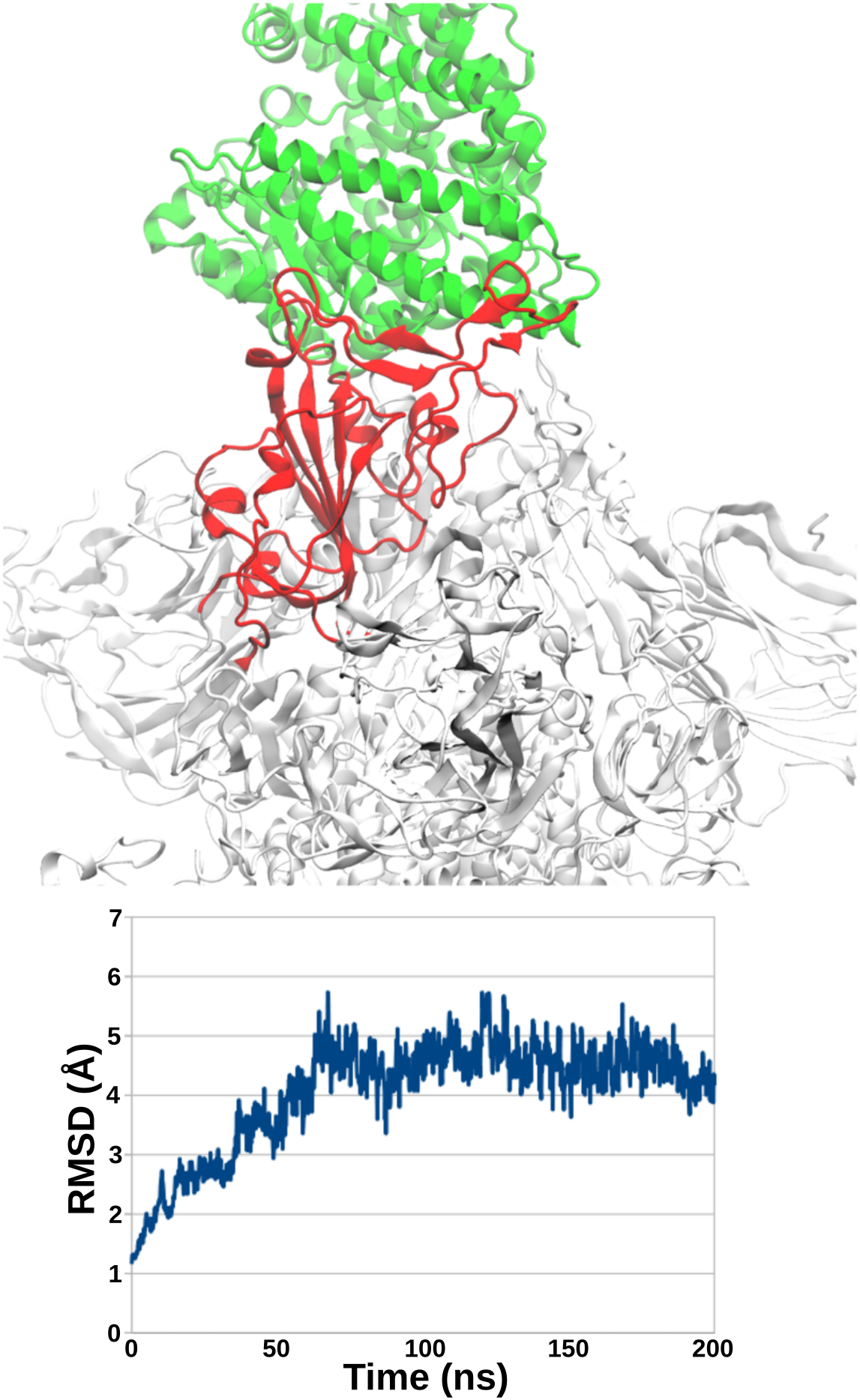
Stability of RaTG13 spike complexed with hACE2. In the upper panel is reported a representative structure of the complex after having reached the equilibrium. ACE2, RBD and spike reminder are depicted in green, red and white, respectively. In the lower panel is reported the RMSD (root mean square deviation) of the entire complex (heavy atoms only) along the 200 ns unbiased MD simulation.

**Figure S6.**
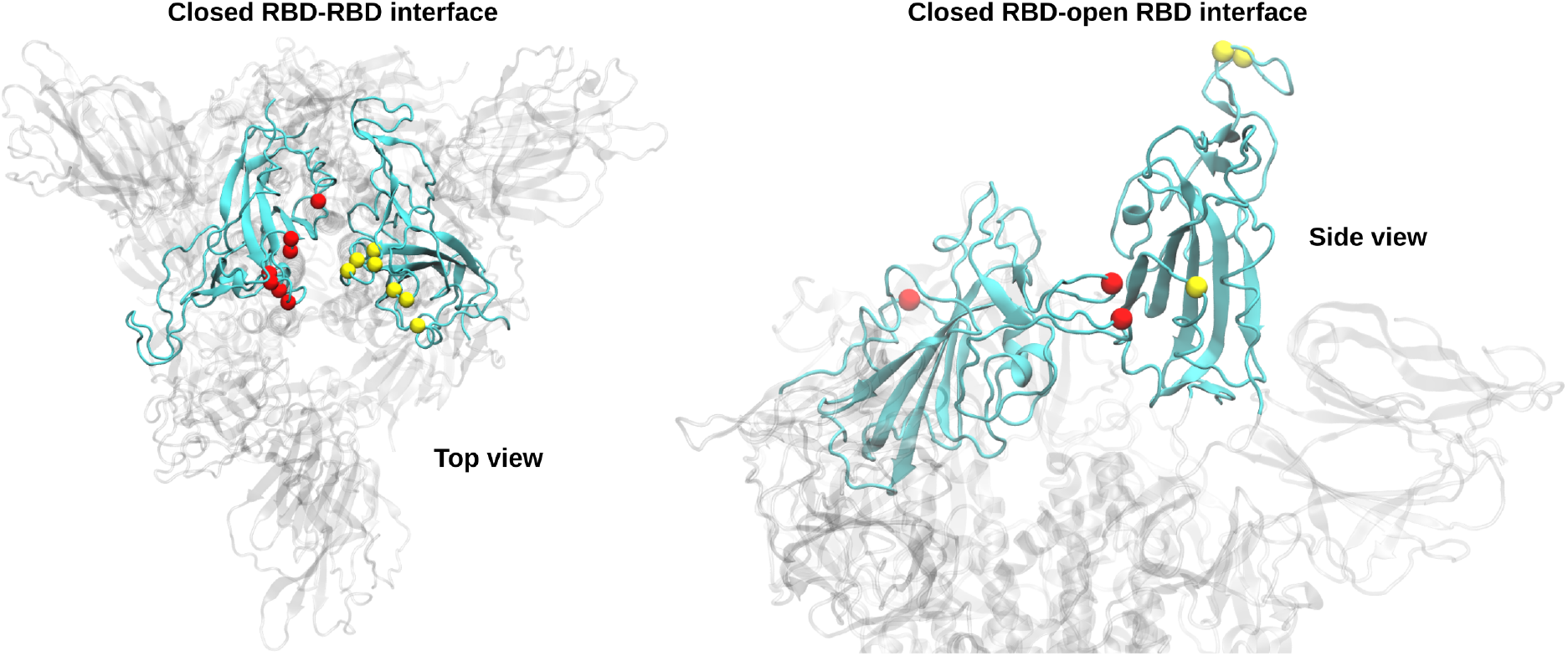
Mutations at the RBD-RBD interface. Mutations from RaTG13 to SARS-CoV-2 lying at the RBD-RBD interface between closed protomers or one closed and one open protomer. The proteins are depicted in ribbon, with interacting RBDs in solid cyan and protein remainder in transparent white. Mutations are depicted in yellow and red spheres depending on the protomer.

**Figure S7.**
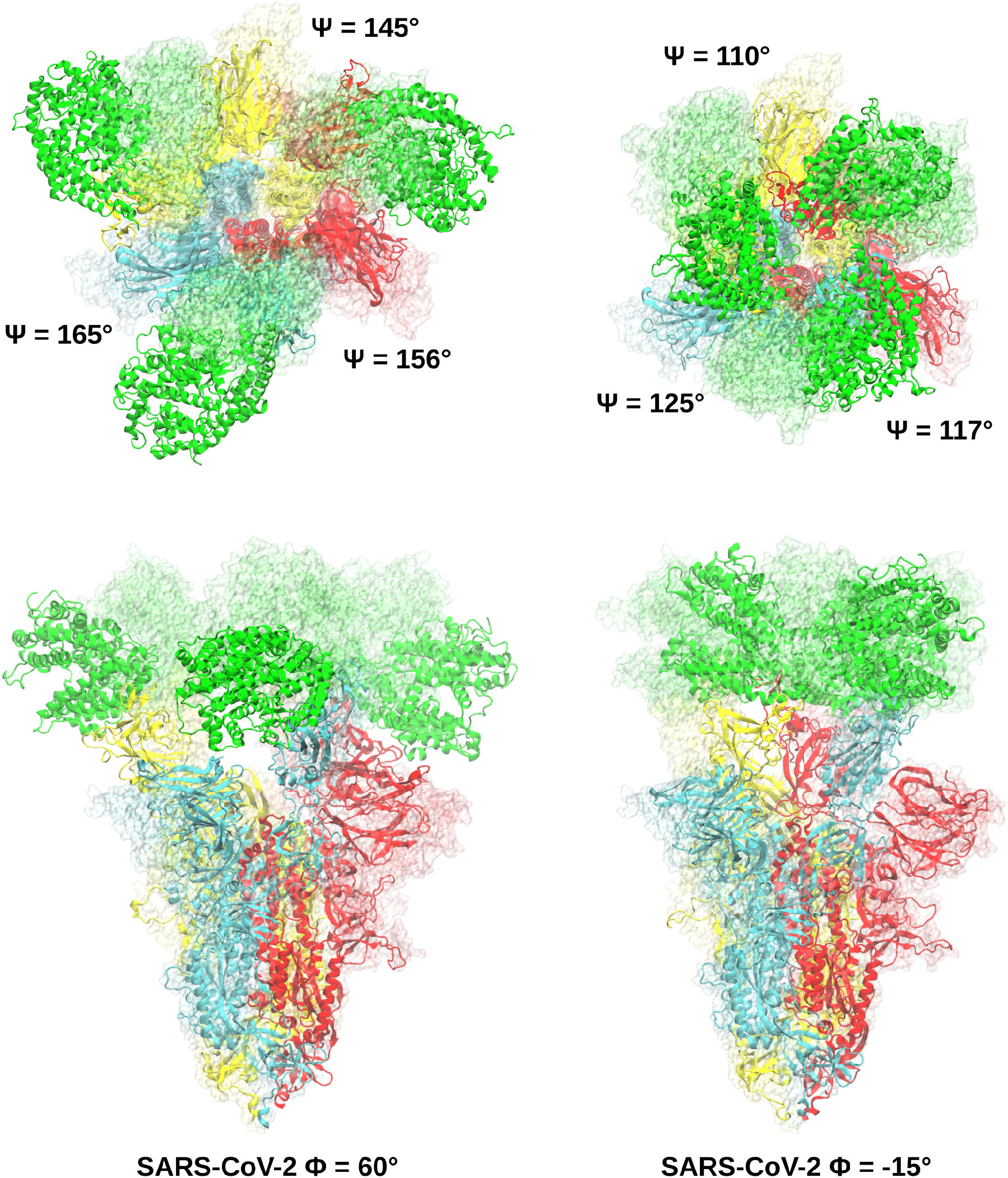
SMD of the 3-up state. Final frames of the SMD simulations at the target Φ angle (60° and −15°). The final Ψ angle necessary to accommodate RBDs rotation is reported for each protomer. The starting conformation is represented in transparent surface, the final conformations in ribbon. The spike protomers are colored in yellow, cyan and red, ACE2 molecules in green.

**Figure S8.**
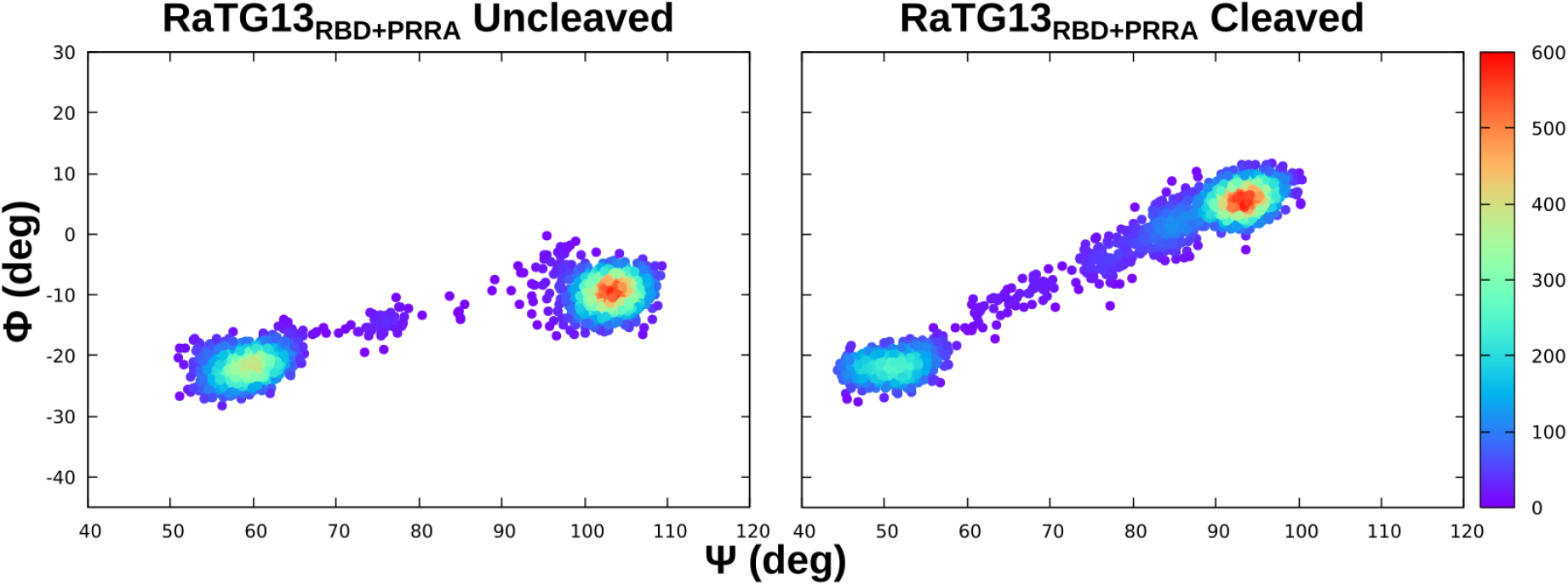
RBD closed-to-1-up transition of RaTG13_RBD+PRRA_ spikes. TMD evolution plotted as a function of density considering the Ψ and Φ angles.

**Figure S9.**
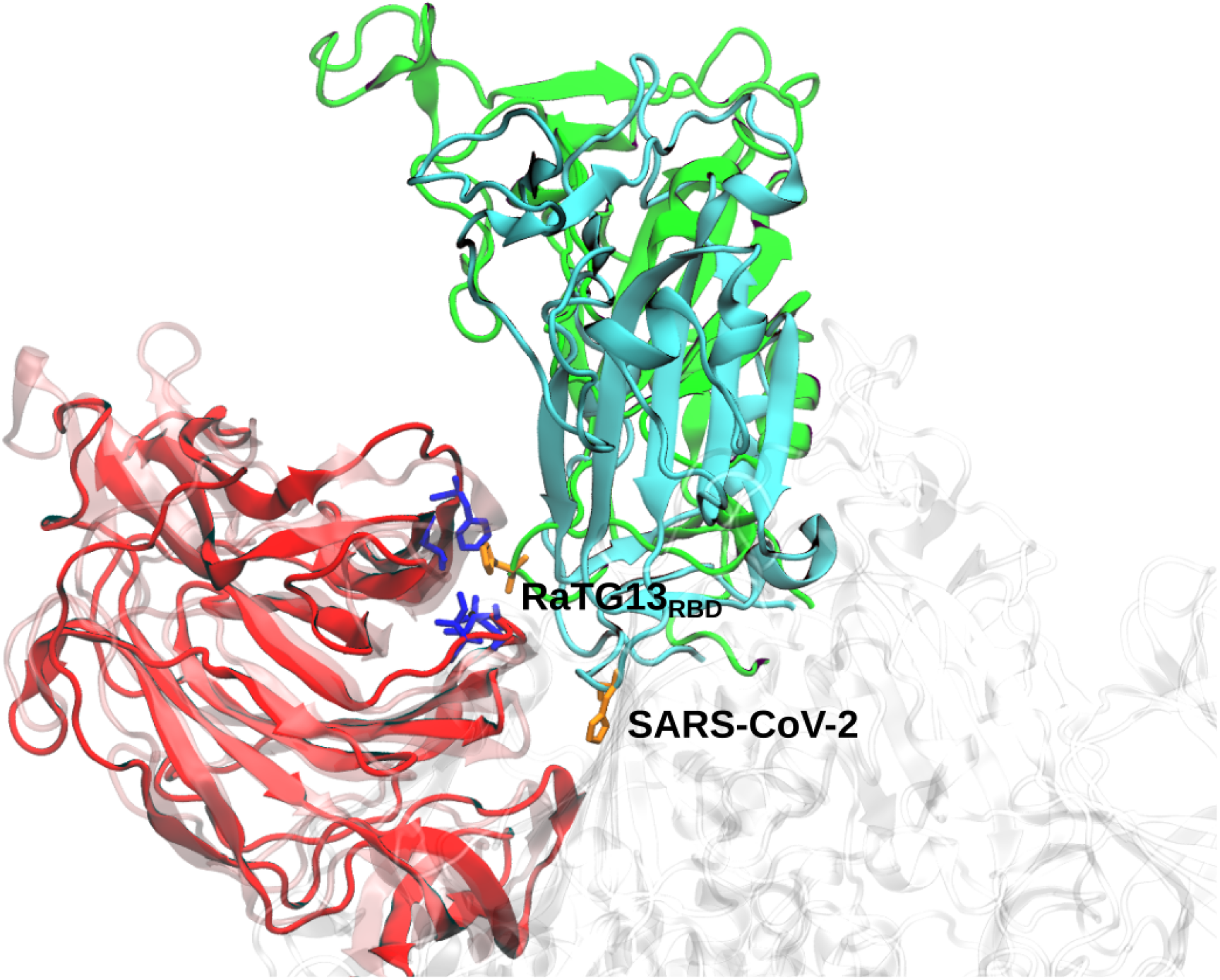
H519 insertion. Representative structure of H519 insertion into the NTD hydrophobic pocket. The NTD, SARS-CoV-2 RBD and RaTG13_RBD_ are depicted in red, cyan and green ribbon, respectively. H519 and the residues forming the hydrophobic pocket are depicted in orange and blue licorice, respectively.

**Figure S10.**
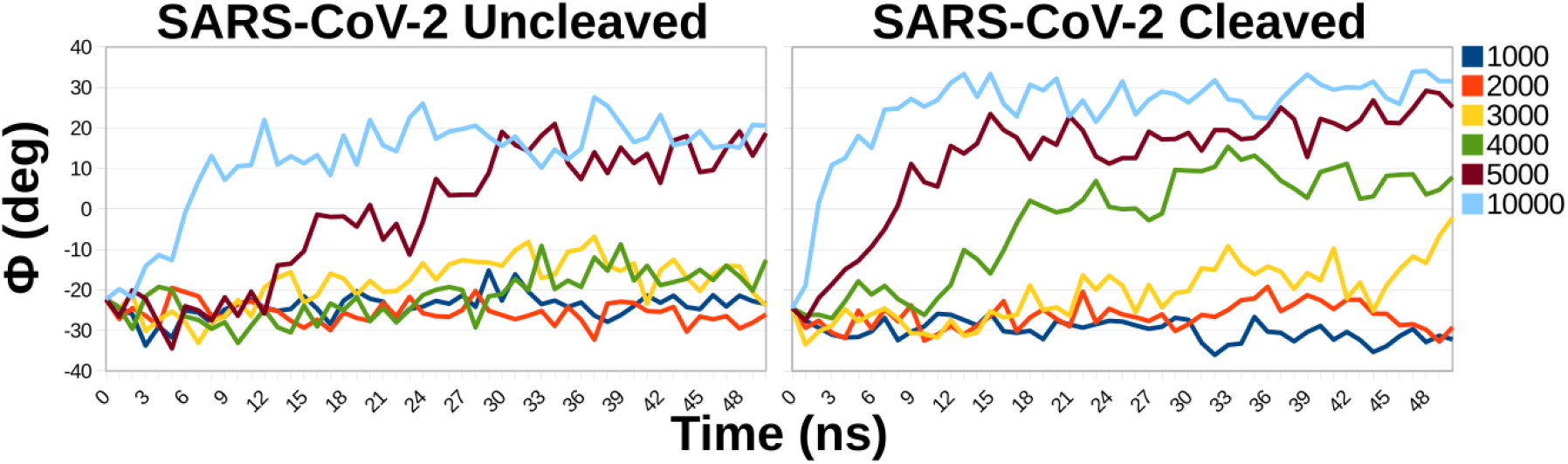
SARS-CoV-2 switch phase TMDs. Short (50 ns) closed-to-1-up TMDs of uncleaved and cleaved SARS-CoV-2 at fixed κ (kJoule/mol/nm). Initial frames were taken from the corresponding complete transition TMDs at the end of the lag phase. The switch was monitored as a function of Φ.

**Table S1.**
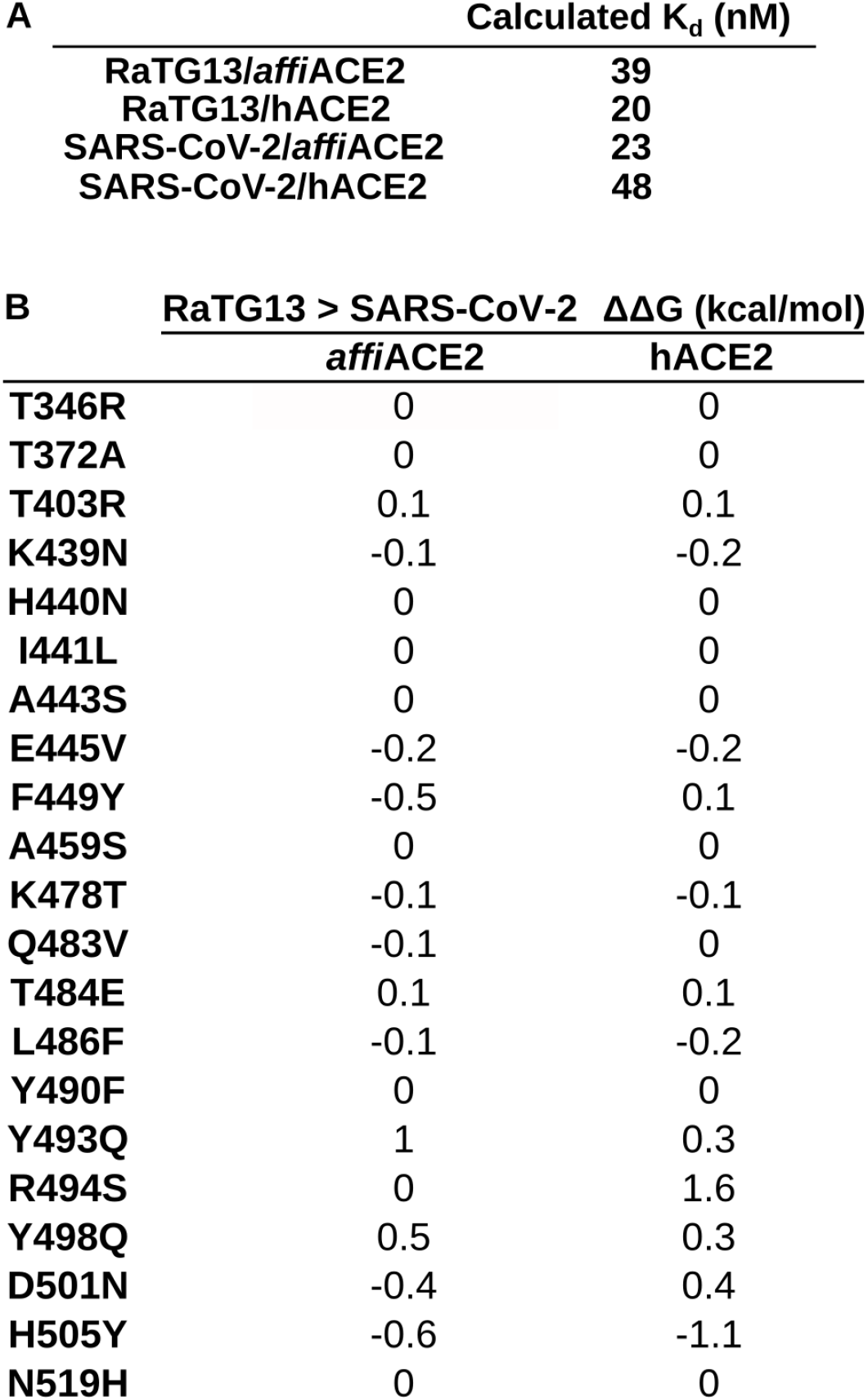
MM-GBSA affinity calculation. (A) Calculated Kd based on free energy calculations. (B) Differential per-residue energy contribution of the RBD mutations to the binding to *affi*ACE2 and hACE2.

**Table S2:** data collection and refinement statistics for the RaTG13 RBD/hACE2 complex datasets.

|  | Crystal Form 1 | Crystal Form 2 |
| --- | --- | --- |
| **Data Collection** |  |  |
| X-ray source | ESRF ID23-EH1 | ESRF ID23-EH1 |
| Processing programs | *XIA2-DIALS, AIMLESS* | *XIA2-DIALS, AIMLESS* |
| Space group | P2<sub>1</sub> | P2<sub>1</sub>2<sub>1</sub>2<sub>1</sub> |
| Cell parameters | a = 83.6 Å; = 115.7°<br>b = 131.2 Å; = 90.0°<br>c = 115.7 Å; = 99.8° | a = 89.9 Å; = 90.0°<br>b = 107.7 Å; = 90.0°<br>c = 251.8 Å; = 90.0° |
| Wavelength (Å) | 0.973 | 0.973 |
| Resolution (Å) | 69.75 – 4.49 (5.02 – 4.49) | 66.42 – 6.50 (7.26 – 6.50) |
| Total reflections | 77828 (22169) | 44159 (12604) |
| Unique reflections | 14753 (4138) | 5180 (1424) |
| *CC1/2*<sup>b</sup> | 0.969 (0.378) | 0.979 (0.531) |
| Redundancy | 5.3 (5.4) | 8.5 (8.9) |
| Mean *I*/*σ*(*I*) | 3.9 (1.2) | 2.8 (0.4) |
| Completeness (%) | 99.4 (98.5) | 99.7 (99.2) |
| *R*<sub>sym</sub><sup>c</sup> | 0.278 (1.178) | 0.459 (2.905) |
| *R*<sub>pim</sub><sup>d</sup> | 0.198 (0.850) | 0.222 (1.407) |
| **Refinement** |  |  |
| *R*<sub>work</sub>/*R*<sub>free</sub><sup>e</sup> | 0.2777/0.2944 | 0.3179/0.3450 |
| Structural refinement protocol | Restrained, Individual B factors + TLS | Rigid Body only |
| Average B-factor (Å<sup>2</sup>) | 191.08 | N/A |
| Total number of atoms | 13101 | 13101 |
| Number of protein atoms | 12791 | 12791 |
| RMS bond lengths (Å) | 0.004 | 0.005 |
| RMS bond angles (°) | 0.82 | 0.81 |
| Ramachandran Favored (%) | 96.1 | 93.1 |
| Ramachandran Allowed (%) | 3.9 | 6.6 |
| Ramachandran Outliers (%) | 0.0 | 0.3 |

<sup>a</sup> Values in parentheses are for reflections in the highest resolution shell.
<sup>b</sup> Resolution limits were determined by applying a cut-off based on the mean intensity correlation coefficient of half-datasets (*CC1/2*) approximately of 0.5 (PA & K, 2012) .
<sup>c</sup> $R_{sym} = [ \sum_{hkl} \sum_j | I_{hkl,j} - \langle I_{hkl} \rangle | ] / [ \sum_{hkl} \sum_j I_{hkl,j} ]$ , where $I_{hkl}$ is the observed intensity for a reflection and $\langle I_{hkl} \rangle$ is the average intensity obtained from multiple observations of symmetry-related reflections.
<sup>d</sup> $R_{pim} = [ \sum_{hkl} (1/(n-1))^{1/2} \sum_j | I_{hkl,j} - \langle I_{hkl} \rangle | ] / [ \sum_{hkl} \sum_j I_{hkl,j} ]$ where $I_{hkl}$ is the observed intensity for a reflection and $\langle I_{hkl} \rangle$ is the average intensity obtained from multiple observations of symmetry-related reflections.
<sup>e</sup> *R*<sub>free</sub> values are calculated based on 5% randomly selected reflections.

